# A flexible cross-correlation based population model of interaural time difference coding in barn owl’s midbrain

**DOI:** 10.64898/2026.04.29.721697

**Authors:** Brian J. Fischer, Ruqhaiya Fatima Syeda, José L. Peña

## Abstract

The cross-correlation model has long served as the standard computational framework for describing interaural time difference (ITD) processing in the barn owl’s auditory system. While successful in explaining initial sinusoidal responses at the site of coincidence detection in the nucleus laminaris, this previous standard model fails to capture the full diversity of ITD tuning observed in the inferior colliculus (IC), where neurons exhibit sharper-than-sinusoidal ITD tuning, nonlinear frequency integration, level-dependent gain control, and interaural level difference (ILD)-dependent modulation of ITD selectivity. Here we present a modified cross-correlation model that addresses these limitations through the addition of parameterized gain control, linear filters with inhibitory surround structure, static nonlinearities, and ILD-dependent modulation of the cross-correlation computation. We show that divisive gain control produces realistic rate-level functions, including non-monotonic responses. Furthermore, inhibitory weights in the linear filter, combined with a threshold or expansive nonlinearity, generate sharper-than-sinusoidal ITD tuning consistent with experimental observations. This model reproduces both linear and nonlinear two-tone frequency integration and demonstrates that independent variation of filter bandwidth and nonlinearity shape accounts for the experimentally observed lack of correlation between side-peak suppression and frequency tuning width across the neuronal population. In addition, ILD-dependent modifications to the model produce shifts in best ITD and reductions in ITD tuning strength, as observed in the lateral shell of the central nucleus of the IC. The model parameters can be efficiently determined using simulation-based inference, enabling generation of realistic neuronal populations. Thus, this flexible, analytically tractable framework provides a foundation for investigating population coding of auditory space in the owl’s midbrain.

## Introduction

Barn owls localize sounds in the horizontal plane using interaural time differences (ITDs), the microsecond-scale delay between a sound’s arrival at each ear (Moiseff and Konishi, 1981; Moiseff, 1989). Models of ITD coding date back to Jeffress (1948), who proposed that a system of axonal delay lines and coincidence-detecting neurons could create a place code for ITD. In barn owls, ITD is initially computed in a specialized circuit comprising axonal delay lines, which introduce precise time delays in neural signals from one ear relative to the other, and coincidence detector neurons in the nucleus laminaris (NL) that are sensitive to the simultaneous arrival of temporally aligned signals. Carr and Konishi (1988) demonstrated that NL afferents exhibit systematic conduction delays that span the owl’s natural ITD range, providing both anatomical and physiological support for a Jeffress-type delay-line circuit model.

The ITD pathway following NL targets the inferior colliculus (IC), a midbrain structure composed of multiple subnuclei, which is a central component of the midbrain and forebrain auditory pathways. In the barn owl’s IC, ITD is processed within and across frequency channels, ultimately generating a map of auditory space in the external nucleus of the inferior colliculus (ICx) (Knudsen and Konishi, 1978). The core of the central nucleus of the inferior colliculus (ICcc) receives direct input from NL and contains neurons sensitive to ITD and narrowly tuned to frequency (Takahashi and Konishi, 1988; Wagner et al., 2002). These neurons often show phase-ambiguous responses to pure tones, responding equally to ITDs that are integer multiples of the tone’s period. The lateral shell of the central nucleus (ICcl) is the first midbrain region where ITD and interaural level difference (ILD) pathways converge following their initial segregation in the brainstem (Adolphs, 1993a). Neurons in ICcl retain sensitivity to ITD and show ILD tuning (Adolphs, 1993b; Fischer et al., 2007). ICx, receiving input from ICcl, contains space-specific neurons that respond selectively to sounds originating from specific directions in auditory space (Knudsen et al., 1977; Knudsen and Konishi, 1978). These neurons are tuned to combinations of ITD and ILD across a broad frequency range, which correspond to specific azimuthal and elevational locations (Moiseff, 1989; Brainard et al., 1992; Euston and Takahashi, 2002).

The barn owl’s auditory system is a well-established model in which computational theories have successfully described the processing of sensory information leading to behavior (Konishi, 1991; Keller and Takahashi, 1996; Peña and Konishi, 2001; Fischer et al., 2009; Fischer and Peña, 2011). One of the most successful computational frameworks is the cross-correlation model of ITD computation. In signal processing terms, the Jeffress-style architecture of delay lines and coincidence detectors performs a cross-correlation of the inputs from the left and right ears. This cross-correlation model analyzes the similarity between the binaural signals as a function of time delay, effectively sliding one signal relative to the other to identify the delay yielding maximal similarity.

Cross-correlation models have been used to describe both the initial computation of ITD in NL and the transformation of ITD information in the IC culminating in space-specific responses in ICx. Numerous studies confirm that NL neurons behave in a manner consistent with cross-correlators. Carr and Konishi (1988) found systematic variations in axonal conduction delays with NL depth, supporting the anatomical substrate of a delay-line model. Physiological recordings further validate the model’s predictions: bilateral inputs to NL exhibit matched spectral tuning (Fischer and Peña, 2009) and responses reflect binaural correlation (Albeck and Konishi, 1995; Fischer et al., 2008). These results support the view that NL neurons act effectively as auditory cross-correlators.

Although the cross-correlation model also accounts for several aspects of ITD processing in the IC, important differences emerge between neural responses and predictions of the cross-correlation model in IC. While many ICcc and ICcl neurons show periodic ITD tuning, inhibition sharpens responses such that tuning curves deviate from pure sinusoids (Fujita and Konishi, 1991). In ICx, frequency integration resolves phase ambiguity seen in upstream areas, and inhibition further shapes responses leading to the emergence of a space map. Several key observations underscore the need for a modification of the standard cross-correlation model to describe IC response: IC responses to stimulus level reflect gain control; ITD tuning to tones is sharper than sinusoidal in ICcl and ICx (Takahashi and Konishi, 1986; Fujita and Konishi, 1991); frequency integration in IC neurons can be linear or nonlinear (Takahashi and Konishi, 1986; Mori, 1997; Peña and Konishi, 2000); side peak suppression (SPS) for broadband stimuli is not directly correlated with frequency tuning width (Mazer, 1998); best ITD and ITD tuning strength can vary with ILD (Fischer et al., 2007). The standard linear cross-correlation model is insufficient to explain this range of response diversity. Previous efforts to address these limitations have introduced frequency-dependent weights and nonlinear transformations (Keller and Takahashi, 2005; Fischer et al., 2009; Gorman et al., 2021).

Because population coding plays a critical role in sound localization (Ferger et al., 2021), a revised model must be capable of producing realistic population responses. In this study, we present a modified cross-correlation framework that accommodates response diversity while retaining analytical tractability. The model’s parameters are interpretable and allow systematic control over frequency and ITD tuning, enabling future exploration of population coding strategies in the owl’s auditory midbrain.

## Methods

### The modified cross-correlation model

The front-end model for the processing of sounds in the barn owl’s auditory system is a modified version of models used in previous studies (Fischer et al., 2009; Beckert et al., 2020). The inputs to the model are the sound inputs to the left and right ears, denoted 𝑠_𝐿_(𝑡) and 𝑠_𝑅_(𝑡), respectively. Input sound signals to the left and right ears are filtered with a bank of gammatone filters that model frequency filtering in the barn owl cochlea (Köppl, 1997)

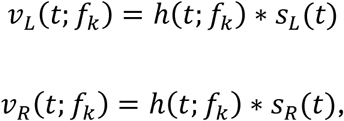

where ∗ denotes convolution and ℎ(𝑡; 𝑓_𝑘_) is the gammatone filter with center frequency 𝑓_𝑘_. The 𝑄_10_ values of the gammatone filters depend on the center frequency of the filter as 𝑄_10_(𝑓) = 0.74√𝑓 to model the responses of auditory nerve fibers in the barn owl (Köppl, 1997).

Operations are performed within each frequency channel created by the gammatone filters to extract ITD- and ILD-dependent cues. Level-dependent information is extracted from the outputs of the gammatone filters on the left and right sides. First, the energy in gammatone filter outputs 𝑣_𝐿_(𝑡; 𝑓_𝑘_), 𝑣_𝑅_(𝑡; 𝑓_𝑘_) is computed by low pass filtering the square of the gammatone filter outputs:

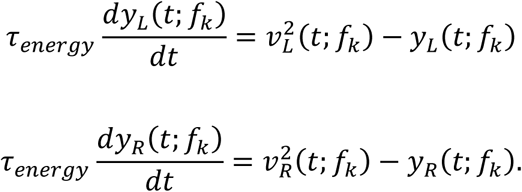

An estimate of the time-dependent ILD is computed from the difference in the logarithm of the energy on the right and left sides as

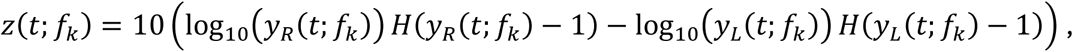

where 𝐻(𝑦) is the unit step function. Including the unit step function of 𝑦 − 1 rectifies the logarithm of the energy to be non-negative. This component of the model is equivalent to the model presented in (Fischer et al., 2009).

ITD-dependent information is extracted from the outputs of the gammatone filters on the left and right sides using a modified cross-correlation model (Albeck and Konishi, 1995; Keller and Takahashi, 1996; Fischer et al., 2008). The cross-correlation model is modified from previous versions (Fischer et al., 2009; Beckert et al., 2020) to allow for neuron-specific dependence of ITD tuning on stimulus level and ILD. First, the gammatone filter outputs are scaled by a measure of their energy to perform a gain control operation:

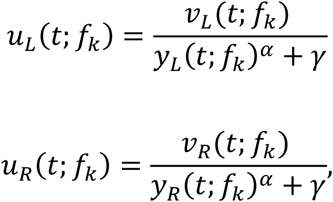

where the exponent 𝛼 and the constant 𝛾 are neuron-specific values that influences the shape and threshold of the rate-level function, respectively. The running cross correlation 𝑥(𝑡, Δ_𝐿_, Δ_𝑅_; 𝑓_𝑘_) is a low-pass filtered version of the square of the sum of the input signals:

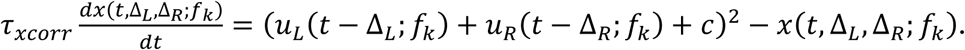

The internal delays Δ_𝐿_, Δ_𝑅_ cover the range 0 – 300 𝜇𝑠 in steps of 5 𝜇𝑠. Using the square of the sum, rather than the product, of the left and right signals allows the model to respond to monaural and binaural inputs (Fischer et al., 2008, 2009).

The cues 𝑧(𝑡; 𝑓_𝑘_) and 𝑥(𝑡, Δ_𝐿_, Δ_𝑅_; 𝑓_𝑘_) are combined in neuron-specific forms to create an input current that drives an adaptive exponential integrate and fire model for IC neurons (Brette and Gerstner, 2005; Fontaine et al., 2014). The input current is formed by first multiplying the running cross-correlation by a matrix that models lateral inhibition between internal delays (Fujita and Konishi, 1991)

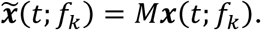

The matrix 𝑀 is a difference of Gaussians function of the internal delay difference Δ_𝐿_− Δ_𝑅_ and the vector 𝒙(𝑡; 𝑓_𝑘_) contains the running cross-correlation across all internal delays

The transformed cross-correlation vector then is processed to model ILD-dependent variation in ITD tuning, as observed at the site of coincidence detection in NL (Peña et al., 1996; Viete et al., 1997). ITD tuning can vary with ILD through a shift in the best ITD or a change in the strength of ITD tuning. To model these changes, the component of the cross-correlation vector, and thus the internal delay that drives the neuron’s response can vary with ILD and the magnitude of the cross-correlation is scaled using a sigmoidal function of ILD. Finally, a weighted sum is computed across frequency channels

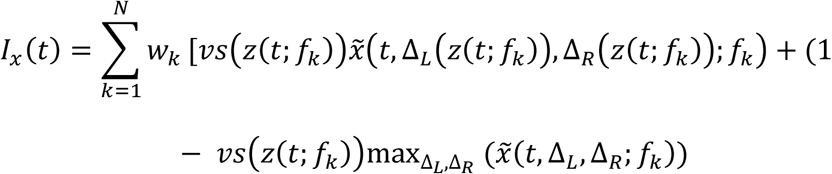

where 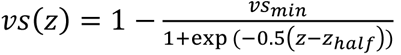.

The input current to the spiking neuron model is formed by processing this input with a rectified power-law static nonlinearity

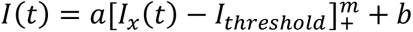. Model spikes are produced using an adaptive exponential integrate and fire neuron (Brette and Gerstner, 2005; Fontaine et al., 2014). The dynamics of the membrane potential 𝑉 and the adaptation current 𝑤 are described by the differential equations

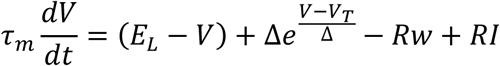

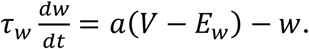

The membrane potential is initialized to the resting potential. The adaptation current is initialized to the steady-state value that would be achieved at the given initial membrane potential:

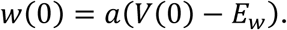

Spikes are produced when the membrane potential exceeds a threshold that depends on the parameter 𝑉_𝑇_ and the voltage derivative (Peña and Konishi, 2002; Fontaine et al., 2014). The threshold is given by 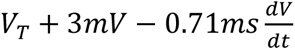.

All differential equations are solved numerically using Euler’s method with a time step of 1/30 ms.

### Parameter determination

A complete characterization of the model requires determining the parameters that control the gain control, the linear filter, the static nonlinearity, and the input scaling to the spiking neuron model. Each model neuron is specified by a set of desired response properties that define the neuron’s tuning characteristics. The desired response properties are: best frequency, frequency tuning half-width, best ITD, side-peak suppression (SPS), ITD shift rate with ILD, ITD peak-trough reduction with ILD, rate-level function shape parameters (𝐴_2_, 𝐴_3_, 𝐴_4_), maximum firing rate, and minimum firing rate. The desired rate-level function shape is specified using the parametric form introduced by Köppl and Yates (1999) for auditory-nerve fiber rate-level functions. This function is defined as

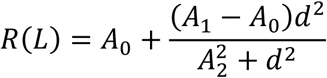

where 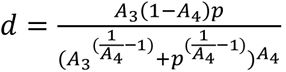, 𝑝 = 10^𝐿/20^, and 𝐿 is the average binaural level (ABL) in dB SPL. 𝐴_0_ is the minimum response, 𝐴_1_is the maximum response, 𝐴_2_controls the half-maximal response level, 𝐴_3_controls the level at which the response transitions from growth to saturation, and 𝐴_4_controls the sharpness of the transition. The parameters best frequency and best ITD directly set the center of the linear filter. The ITD shift rate directly sets the rate at which the linear filter center shifts along the ITD axis as a function of ILD. The ITD peak-trough reduction parameter and its half-maximal ILD directly set the parameters of the sigmoidal function that modulates ITD tuning strength with ILD.

Several model parameters can be computed directly from the desired response properties, while others require a fitting procedure. The parameter determination proceeds in three stages: (1) fitting the gain control parameters using simulation-based inference, (2) fitting the linear filter bandwidth and nonlinearity power using simulation-based inference, and (3) computing the gain and bias of the input to the spiking neuron analytically.

Stage 1: Gain control parameters. The gain control exponent α and constant γ determine the shape of the rate-level function of the cross-correlation output prior to the linear filter and nonlinearity. These parameters are determined using simulation-based inference (SBI) with sequential neural posterior estimation (SNPE) (Cranmer et al., 2020). A uniform prior distribution is defined over the ranges of α and γ. For each of N samples drawn from the prior, the model computes the rate-level function of the linear cross-correlation output for broadband noise at the neuron’s best ITD across ABL values from 0 to 80 dB SPL. Each simulated rate-level function is fit with the Köppl and Yates (1999) parametric form, yielding summary statistics (𝐴_2_, 𝐴_3_, 𝐴_4_). A mixture density network (MDN) is trained on the parameter summary statistic pairs to approximate the posterior distribution 𝑝(𝛼, 𝛾|𝐴_2_, 𝐴_3_, 𝐴_4_). The maximum a posteriori (MAP) estimate from this posterior, conditioned on the desired 𝐴_2_, 𝐴_3_ and 𝐴_4_ values, provides the gain control parameters for the neuron.

Stage 2: Filter bandwidth and nonlinearity power. The frequency bandwidth of the linear filter (𝜎_𝑓_) and the power of the power-law static nonlinearity (𝑚) jointly determine the neuron’s frequency tuning width and side-peak suppression for broadband noise. Before fitting these parameters, the threshold of the static nonlinearity 𝐼_𝑡ℎ𝑟𝑒𝑠ℎ𝑜𝑙𝑑_ is set to the minimum of the linear cross-correlation response across ABL, ensuring that the nonlinearity rectifies at the appropriate operating point. SBI with SNPE is again used to approximate the posterior distribution over 𝜎_𝑓_and 𝑚. For each sample drawn from a uniform prior over these two parameters, the model computes the ITD tuning curve and frequency tuning curve for broadband noise at the neuron’s best ITD. Summary statistics are extracted from these responses: SPS from the ITD tuning curve and frequency half-width from the frequency tuning curve. An MDN is trained to approximate the posterior 𝑝(𝜎_𝑓_, 𝑚 | SPS, frequency half-width). Parameter estimates are obtained by repeatedly sampling from the posterior conditioned on the desired SPS and frequency half-width values, evaluating each sample, and accepting the first sample whose SPS and frequency half-width fall within tolerance (1 dB for SPS, 0.25 kHz for frequency half-width) of the desired values, up to a maximum of 100 iterations.

Stage 3: Input scaling. The gain (a) and bias (b) that scale the input current to the spiking neuron model are computed analytically to map the model’s output range to the desired firing rate range. The f-I curve of the adaptive exponential integrate-and-fire neuron is computed numerically by simulating responses to constant input currents across a range of values, and its inverse 𝑓^−1^is obtained by interpolation. The cross-correlation model response is then evaluated at two stimulus conditions: a high-level broadband noise stimulus (80 dB ABL) at the neuron’s best ITD, producing the maximum response of the static nonlinearity component 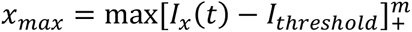, and a low-level stimulus (0 dB ABL), producing the minimum response 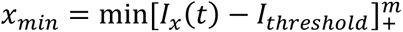. The gain and bias are determined by solving the linear system

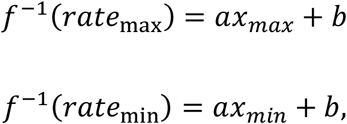

where 𝑟𝑎𝑡𝑒_max_ and 𝑟𝑎𝑡𝑒_min_ are the desired maximum and minimum firing rates. This ensures that the spiking neuron produces the desired firing rate range for the range of cross-correlation model outputs.

For the SBI fitting in both Stage 1 and Stage 2, N = 5,000 samples were drawn from the prior distribution. The MDN was trained using the SNPE algorithm as implemented in the sbi Python package (Tejero-Cantero et al., 2020). Simulations producing invalid summary statistics (NaN values) were excluded from training. The three-stage procedure is applied independently to each neuron in the population, with stages executed sequentially because the gain control parameters from Stage 1 influence the responses used in Stage 2, and the filter and nonlinearity parameters from Stage 2 determine the response range used in Stage 3.

See https://github.com/brian-fischer/Flexible-xcorr-model for the model code.

## Results

We aimed to develop a model that can account for the diverse ITD tuning properties observed in the barn owl’s ICcl and ICx, including their dependence on stimulus frequency bandwidth and ILD. Here we present a modification to the standard cross-correlation model of ITD computation that includes parameterized linear filters and nonlinearities that can be fit to produce the ITD tuning observed in ICcl and ICx. We define the *standard cross-correlation model* (Figure 1A) as follows: input sound signals are filtered through a bank of bandpass filters (here modeled as gammatone filters), and a running cross-correlation is computed in each frequency channel. The cross-correlation outputs are then summed across all channels, and the response is time-averaged. The standard cross-correlation model was modified to include gain control for stimulus amplitude dependence (Peña et al., 1996; Köppl and Yates, 1999), ILD-dependence of the cross-correlation computation (Viete et al., 1997), nonuniform weighting across frequency and ITD (Gold and Knudsen, 2000; Cazettes et al., 2014), and static point nonlinearities to model transformations within the ITD pathway and the spiking nonlinearity (Fujita and Konishi, 1991; Albeck and Konishi, 1995). We first demonstrate the responses of the standard cross-correlation model as a baseline, then show how the modifications to the model allow it to produce different aspects of experimentally observed ITD tuning.

**Figure 1:**
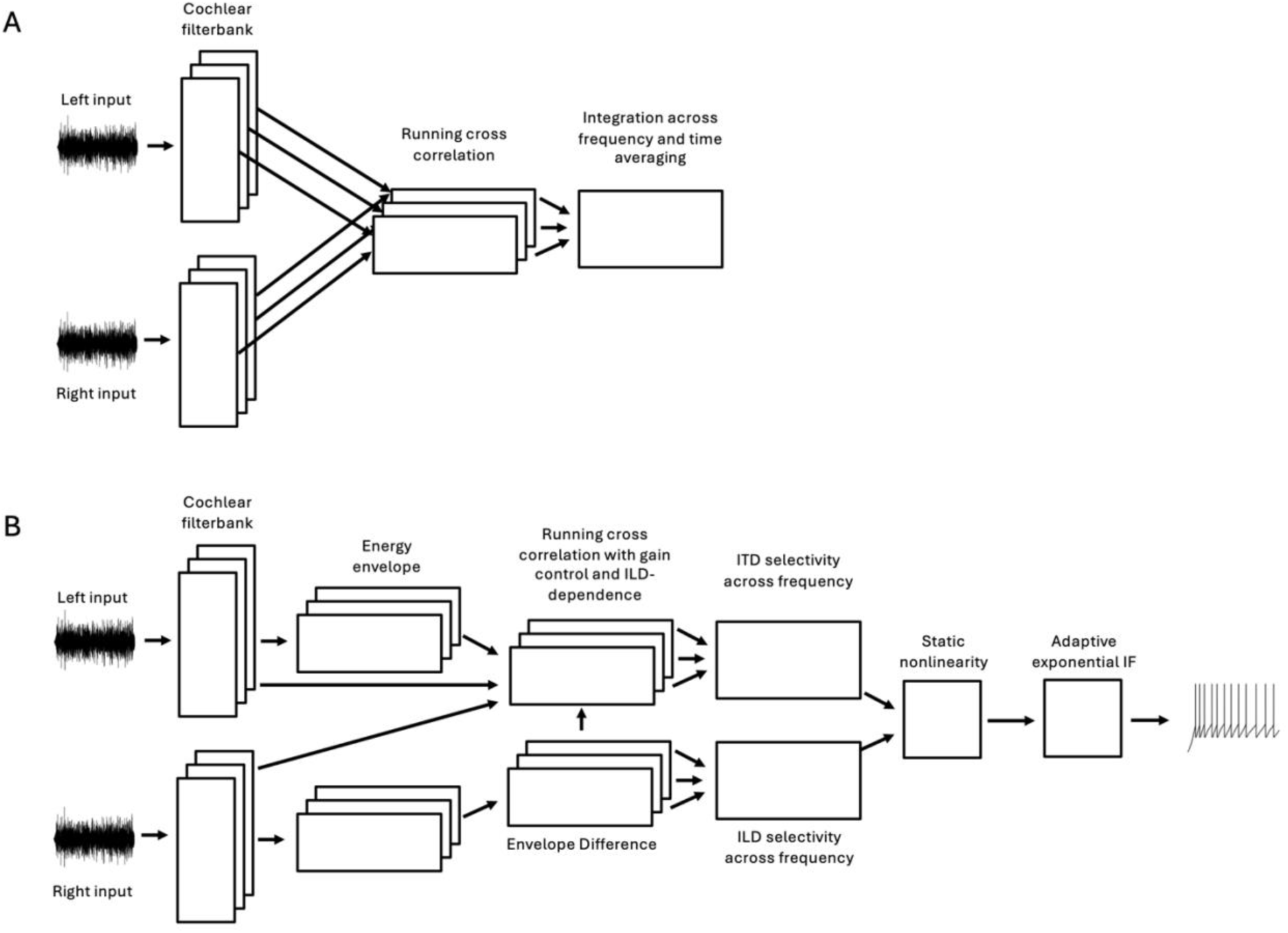
Standard and modified cross-correlation models. (A) The standard cross-correlation model. Input sound signals to the left and right ears are filtered by a bank of gammatone filters modeling cochlear frequency selectivity. A running cross-correlation is computed within each frequency channel, and the outputs are summed across channels and time-averaged to produce an ITD tuning curve. (B) The modified cross-correlation model. The model incorporates divisive gain control at the input to the cross-correlation, a linear filter with neuron-specific weights across frequency and ITD channels (including inhibitory surround structure), ILD-dependent modulation of the cross-correlation output, a static nonlinearity, and an adaptive exponential integrate-and-fire spiking neuron. The ILD pathway (bottom) extracts interaural level differences from the energy envelopes of the gammatone filter outputs. The ITD and ILD pathways converge at the input to the spiking neuron model.

### The standard cross-correlation model

The standard cross-correlation model is based on computing the running cross-correlation within each frequency channel created by a bank of bandpass filters, summing the results across frequencies, and averaging over time.

The standard cross-correlation model produces phase-ambiguous responses to tonal stimuli, resulting in sinusoidal ITD tuning curves with half-widths equal to half the period of the tone (Figure 2A). Some ICcl and ICx neurons exhibit sinusoidal ITD tuning curves for tones, with half-widths closely matching theoretical predictions for the standard cross correlation model (Fujita and Konishi, 1991; Mori, 1997). However, most ICcl and ICx neurons have sharper than sinusoidal ITD tuning curves for tones (Takahashi and Konishi, 1986; Fujita and Konishi, 1991).

**Figure 2:**
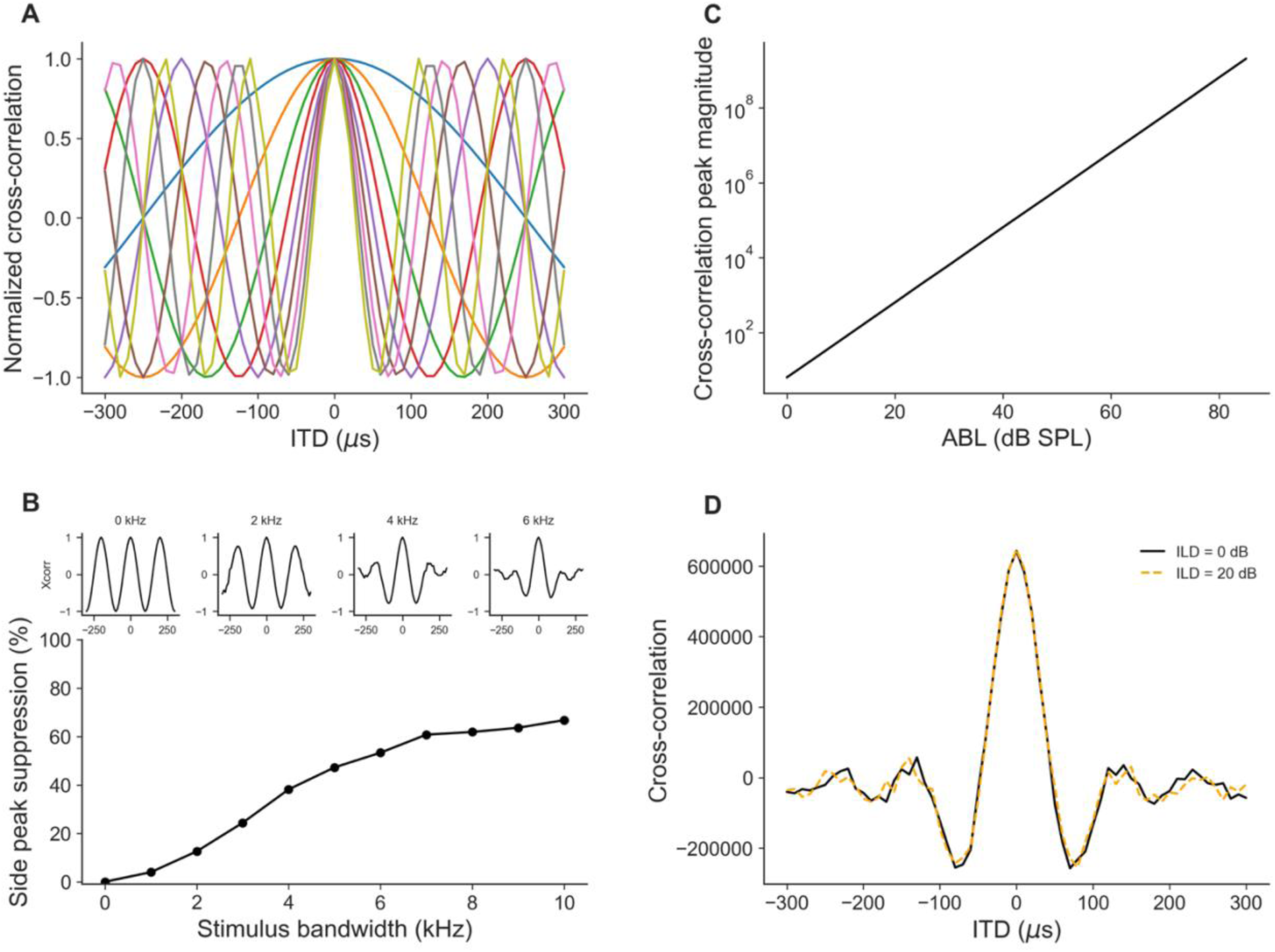
Responses of the standard cross-correlation model. (A) Normalized ITD tuning curves for tonal stimuli at frequencies ranging from 1 to 9 kHz, showing sinusoidal tuning with period equal to the inverse of the stimulus frequency. (B) Side-peak suppression (SPS) as a function of stimulus bandwidth for broadband noise centered at the best frequency. Insets show ITD tuning curves at selected bandwidths. SPS increases sigmoidally with bandwidth. (C) Cross-correlation peak magnitude as a function of average binaural level (ABL) on a log scale, showing exponential growth with stimulus level. (D) ITD tuning curves for two ILD conditions (0 dB and 20 dB). The standard model produces no change in ITD tuning shape or best ITD with ILD.

Frequency integration in ICx is a critical computation for resolving the stimulus ITD from among phase equivalent ITDs (Mazer, 1998; Peña and Konishi, 2000). The standard cross-correlation model uses linear frequency integration. Some ICcl and ICx neurons exhibit ITD tuning consistent with linear integration across frequency, however this occurs for a minority of neurons (Takahashi and Konishi, 1986; Mori, 1997). The standard model with linear frequency integration over the full range of frequency channels predicts that side-peak suppression (SPS) increases with stimulus bandwidth in a sigmoidal manner (Figure 2B). While this general trend is observed in IC neurons, the increase in SPS predicted by the standard cross-correlation model exceeds what is observed in most neurons experimentally (Mazer, 1998) and does not produce diverse SPS behaviors across neurons.

The response magnitude of the standard cross-correlation model grows exponentially with stimulus level (Figure 2C). This contrasts with the sigmoidal increase in response rates as a function of stimulus level observed for most neurons in the auditory system (Peña et al., 1996; Köppl and Yates, 1999; Arthur, 2004). In addition, some IC neurons have non-monotonic responses to stimulus level where the response decreases at high stimulus levels (Knudsen and Konishi, 1978; Arthur, 2004).

Finally, the standard cross-correlation model does not produce changes in ITD tuning shape or best ITD with ILD (Figure 2D). In contrast, ICcl neurons can exhibit weakening ITD tuning or shifts in best ITD with ILD (Fischer et al., 2007), similar to that observed in NL (Viete et al., 1997).

In the following, we present a modified version of the cross-correlation model that removes the inconsistencies of the standard model with the observed responses and allows for flexible matching of model responses to desired responses (Figure 1B).

### Gain control and stimulus level

Beginning in the auditory nerve, auditory neurons respond to stimulus level with approximately sigmoidal increases in firing rate as level increases on a dB scale (Peña et al., 1996; Köppl and Yates, 1999; Arthur, 2004). Previous models have incorporated a gain control operation in the cross-correlation model to influence responses to stimulus level (Peña et al., 1996; Fischer et al., 2009). Here, we include a gain control operation at the input to the running cross-correlation that is parameterized to allow the model to produce rate-level functions with a diversity of experimentally observed shapes, including non-monotonic responses (Figure 3A). The input to the cross-correlation is given by the terms from the left and right:

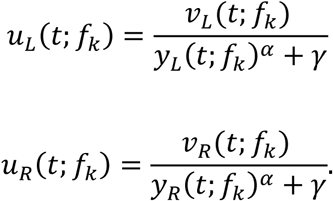

**Figure 3:**
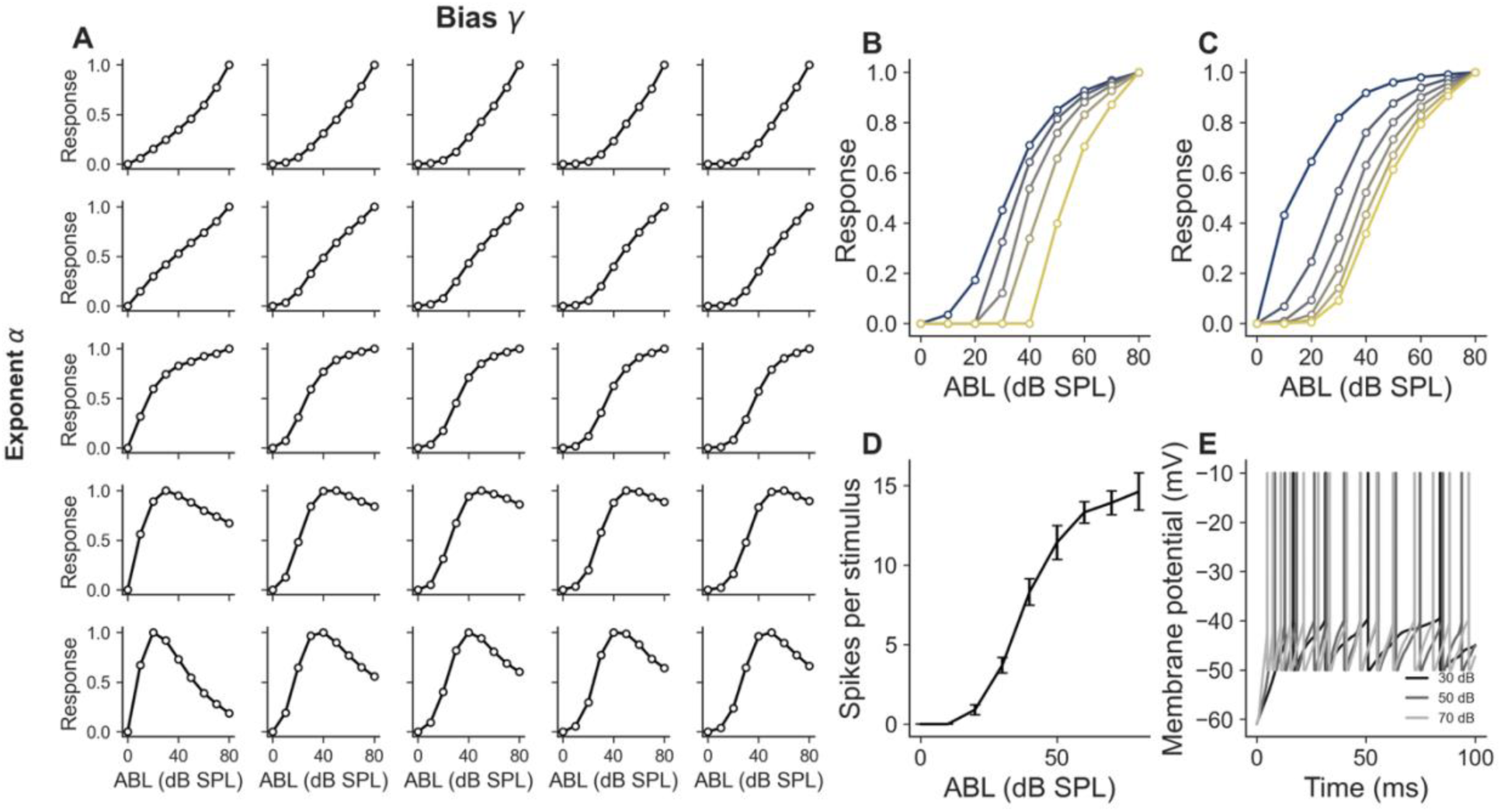
Gain control and responses to stimulus level. (A) Rate-level functions of the modified cross-correlation model for a grid of gain control exponent 𝛼 (columns, 0.45 ≤ 𝛼 ≤ 0.55) and constant bias 𝛾 (rows, 0.5 ≤ 𝛾 ≤ 10) values. Increasing 𝛼 produces a transition from expansive (top, 𝛼 < 0.5) to sigmoidal (center, 𝛼 = 0.5) to non-monotonic (bottom, 𝛼 > 0.5) rate-level functions. Increasing 𝛾 shifts the rate-level function to higher ABL values. (B) Families of rate-level functions showing the effect of varying the threshold of the static nonlinearity (0 – 0.75) for a fixed gain control configuration. (C) Same as B but with a different powers of the static nonlinearity (0.25 – 3). (D) Rate-level function of the spiking neuron model. Error bars indicate standard deviation across stimulus repetitions. (E) Membrane potential traces at ABL values ranging from 30 to 70 dB SPL, showing increasing spike rates with stimulus level while maintaining sensitivity to temporal envelope features across sound levels.

where 𝑣_𝐿_, 𝑣_𝑅_ are the outputs of the cochlear bandpass filters on the left and right sides, respectively, and 𝑦_𝐿_, 𝑦_𝑅_are smoothed versions of the square of the gammatone filter outputs 𝑣_𝐿_, 𝑣_𝑅_. The exponent 𝛼 and the constant 𝛾 are neuron-specific values that influence the curvature and threshold of the rate-level function, respectively (Figure 3A). The model produces rate-level functions that are expansive or approximately linear for exponent values 𝛼 < 0.5, sigmoidal for 𝛼 = 0.5, and non-monotonic for 𝛼 > 0.5. This pattern results because the input-dependent term in the denominator 𝑦(𝑡; 𝑓_𝑘_) is a smoothed version of the square of the gammatone filter output 𝑣(𝑡; 𝑓_𝑘_). Therefore, when 𝛼 < 0.5, the denominator grows slower with ABL than does the numerator, and the rate-level function can be expansive. Conversely, when 𝛼 > 0.5 the denominator grows faster with ABL than does the numerator, and the rate-level function can be non-monotonic where the response decreases at high ABL. Changing the constant 𝛾 produces a horizontal shift of the rate-level function (Figure 3A). A larger constant reduces the relative impact of the input-dependent term in the denominator 𝑦(𝑡; 𝑓_𝑘_), resulting in a shift to higher ABL values before the gain control decreases the response.

The static nonlinearity that models nonlinear processing prior to spiking in ICcl can also influence the rate-level function; the interaction between gain control and the threshold and power of the nonlinearity determines the overall shape of the rate-level response. The static nonlinearity influences the threshold and the saturation in the rate-level function (Figure 3B,C). While the static nonlinearity does influence the rate-level function, it is important to use a divisive gain control operation, as opposed to a static suppressive nonlinearity, to produce the observed dependence of sound level on a dB scale because relying on a static suppressive nonlinearity alone would limit sensitivity to the stimulus envelope. Divisive gain control allows the model to respond to structure in the stimulus envelope as the amplitude of the input signal ranges over several orders of magnitude (Figure 3D,E). Voltage traces of the spiking neuron model at ABL values ranging from 30 to 70 dB show increasing spike rates with stimulus level while maintaining sensitivity of spike timing to temporal features in the input across sound levels. This combination of gain control and spiking nonlinearity produces realistic rate-level functions that match experimental observations of IC neurons.

### ITD tuning widths

Neurons in ICcl and ICx have a range of ITD tuning widths for tonal stimuli, reflecting sharper than sinusoidal ITD tuning in most neurons (Fujita and Konishi, 1991). Sharper than sinusoidal ITD tuning can be produced in the model by a combination of inhibition and a static nonlinearity (Figure 4). Inhibition is modeled using negative weights in the linear filter applied to the cross-correlation output, which forms the input to the static nonlinearity (Figure 4A). In this example, the weights are given by a difference of Gaussian functions. A rectified linear static nonlinearity with a threshold, combined with negative weights in the linear filter, produces sharper than sinusoidal ITD tuning (Figure 4B,C). As observed experimentally, when inhibition is blocked, the input is above the threshold and ITD tuning is approximately sinusoidal, following the input current (Figure 4B,C,D) (Fujita and Konishi, 1991).

**Figure 4:**
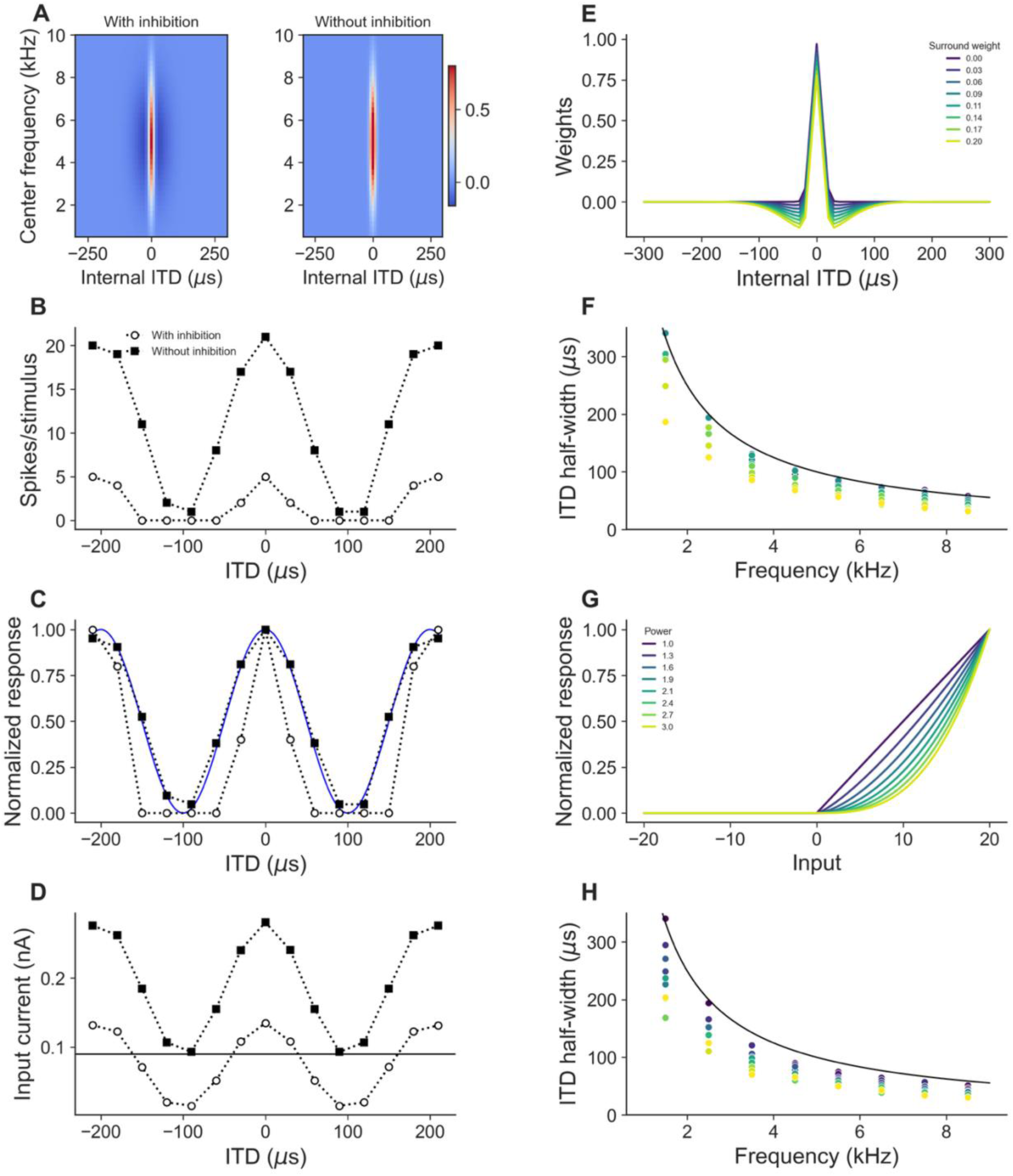
Sharper-than-sinusoidal ITD tuning produced by inhibition and nonlinearity. (A) Linear filter weight matrices with (left) and without (right) inhibitory surround weights, displayed as a function of internal ITD and frequency channel index. The filter is modeled as a difference of Gaussians. (B) ITD tuning curves from the neuron model with surround inhibition (open circles) and without surround inhibition (filled squares). Inhibition sharpens ITD tuning and reduces side-peak responses. (C) Normalized ITD tuning curves from B overlaid with sinusoidal fits (blue curves), showing that the model with inhibition produces sharper-than-sinusoidal tuning. (D) Input current to the spiking neuron as a function of ITD for the two conditions. Horizontal line indicates the spiking threshold. (E) Cross-section of the linear filter weights for different surround inhibition strengths (color-coded, 0.00–0.20). (F) ITD half-width as a function of stimulus frequency for different surround inhibition strengths (color-coded, 0.00–0.20). Black curve shows the sinusoidal prediction (half-period). Increasing surround weight produces systematically sharper ITD tuning. (G) Family of static nonlinearities with powers ranging from 1.0 (linear, yellow) to 3.0 (expansive, dark blue). (H) ITD half-width as a function of stimulus frequency for different nonlinearity powers (color-coded, 1.2–2.8). Increasing nonlinearity power produces sharper ITD tuning independent of the inhibitory mechanism.

The half-width of the ITD tuning curve is directly related to the strength of the inhibitory component of the weights (Figure 4E,F). Figure 4F shows that as surround weight increases from 0.0 to 0.2 across neurons with a rectified linear nonlinearity, the ITD half-width decreases systematically below the sinusoidal prediction (half-period) line. This demonstrates that inhibition and a threshold can sharpen ITD tuning beyond what the standard cross-correlation model predicts.

An alternative mechanism for producing sharper ITD tuning is an expansive static nonlinearity that could result from spiking or dendritic nonlinearities prior to ICcl (Figure 4G). Figure 4H shows that with the power of the nonlinearity varied from 1.0 to 3.0, the ITD half-width decreases systematically with increasing power. Both mechanisms, inhibitory surround weights followed by a threshold and expansive nonlinearity, can independently produce sharper-than-sinusoidal ITD tuning. The flexibility of the model allows for combinations of these mechanisms to capture the diversity of ITD tuning widths observed across the neuronal population.

### Frequency integration and coding of auditory space

A major processing step in the IC is frequency convergence, which enables the computation of auditory space selectivity. Neurons in ICx receive convergent input across multiple frequency channels, and their ITD selectivity depends on whether frequency convergence is linear or nonlinear (Wagner et al., 1987; Mazer, 1998; Peña and Konishi, 2000).

### Linear two-tone frequency integration

A subset of ICx neurons shows ITD tuning for tone combinations that is consistent with linear frequency integration (Takahashi and Konishi, 1986; Mori, 1997; Gorman et al., 2021). Linear frequency integration can be produced by the model when the tone frequencies are far enough apart to avoid nonlinear combination in the gain control mechanism and when the transformation from subthreshold to spiking response is linear (Figure 5A-C). Figure 5A illustrates this for two tones with frequencies 4 and 6 kHz, frequencies far enough apart that they are processed by non-overlapping frequency channels. The response of the model to the combination closely matches the sum of the individual tone responses (Figure 5B). This linear combination is achieved because the static nonlinearity is linear over the relevant input range (Figure 5C).

**Figure 5:**
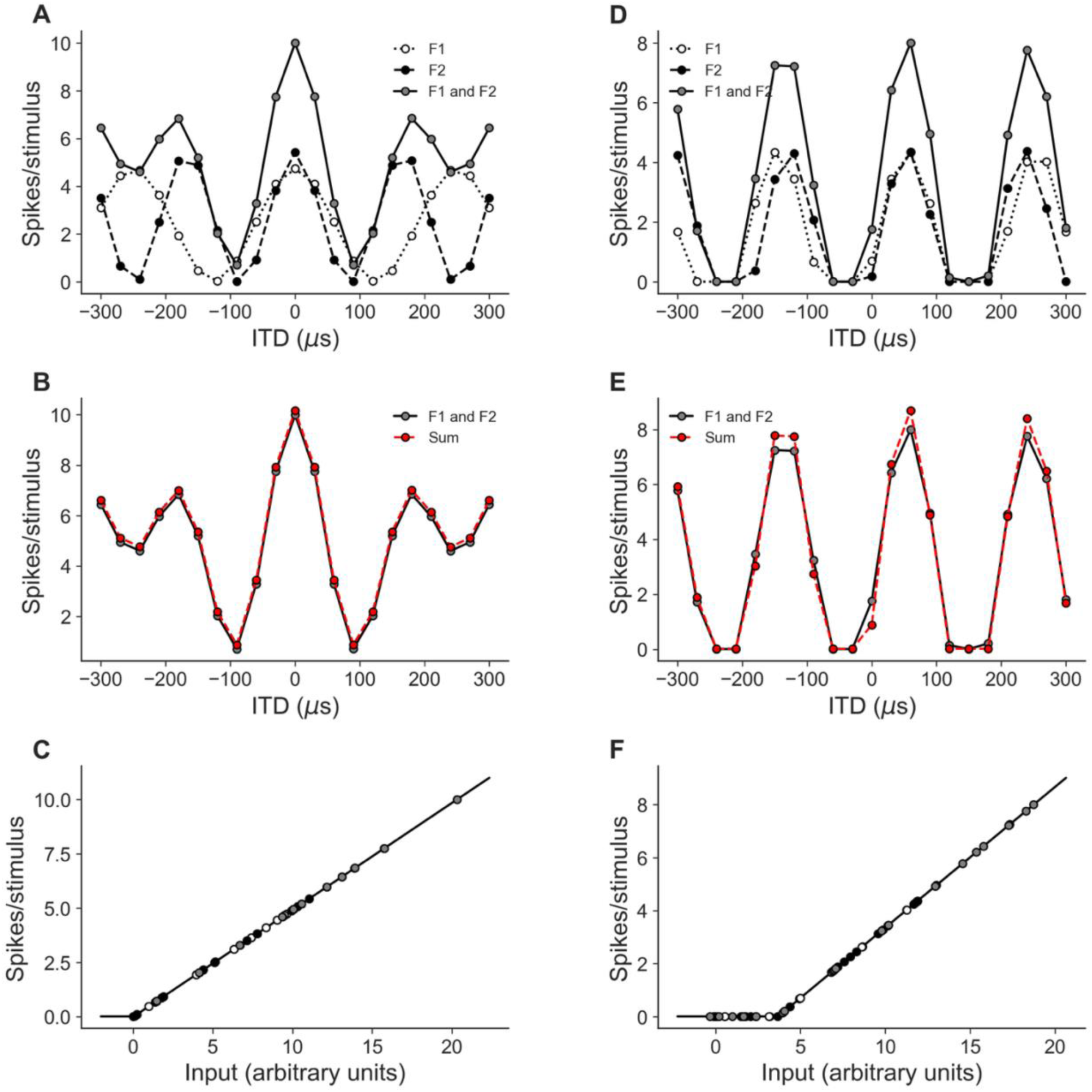
Linear two-tone frequency integration. (A-C) Linear integration of two widely spaced tones (4 and 6 kHz). (A) ITD tuning curves for each tone presented alone (F1, open circles; F2, filled circles) and together (F1 and F2, gray circles). (B) Comparison of the response to the two-tone stimulus (F1 and F2, black) with the sum of the individual tone responses (Sum, red). The close overlap indicates linear frequency integration. (C) Input-output function of the spiking neuron, showing a linear operating regime over the relevant input range. (D-F) Linear integration of two closely spaced tones with a rectified-linear spiking nonlinearity (5 and 5.5 kHz). (D) ITD tuning curves for F1, F2, and F1 and F2. (E) Comparison of F1 andF2 response with the sum, showing approximately linear combination despite the closer frequency spacing. (F) Input-output function showing a rectified-linear nonlinearity with a threshold; linear integration occurs because the alignment of troughs in the individual tone responses minimizes the impact of the threshold.

Additionally, linear frequency integration can also occur for more closely spaced frequencies under appropriate conditions. Mori (1997) shows an example neuron with approximately linear two-tone combination for a condition that might be expected to produce nonlinear responses because the frequencies are close enough (0.5 kHz) to interact through the gain control within each frequency channel (see Figure 1 in (Mori, 1997)). These responses can be reproduced by the model using a rectified-linear static nonlinearity (Figure 5D-F). The response to the two-tone stimulus is approximately linear because the alignment of the troughs of the individual tone ITD curves minimizes the impact of the threshold on the linearity of frequency combination because the two inputs are below threshold for similar ITDs.

### Nonlinear two-tone frequency integration

In contrast to linear frequency integration, many IC neurons show nonlinear frequency integration. Additionally, experimental observations of two-tone frequency integration in ICx neurons show that the form of the combination can be different at the main peak and side peaks of the ITD curve (Takahashi and Konishi, 1986; Mori, 1997). Often, the response to a tone combination at the side peak of the ITD curve is less than the sum of individual responses or the response at the main peak is larger than the sum of individual responses (Takahashi and Konishi, 1986; Mori, 1997).

Nonlinear frequency integration can be produced by the model with many forms of static nonlinearity. Different combinations of the threshold and power of the nonlinearity, together with the alignment of side peaks and troughs of responses to the two frequencies can produce different amounts of facilitation at the main peak and suppression at the side peaks.

Figure 6 demonstrates how inhibition in the linear filter followed by a rectification produces nonlinear frequency integration. The response to two tones at 6 and 8 kHz show side peak suppression and main peak facilitation relative to the sum of individual responses (Figure 6A-D). When the inhibitory weights are removed (Figure 6E), the combination becomes approximately linear (Figure 6F-H). This is consistent with the experimental observations in ICx that blocking GABAergic inhibition leads to linear frequency integration (Mori, 1997). This demonstrates that in the model, the structure of the linear filter, specifically negative weights representing inhibition, plays a critical role in determining whether frequency integration is linear or nonlinear.

**Figure 6:**
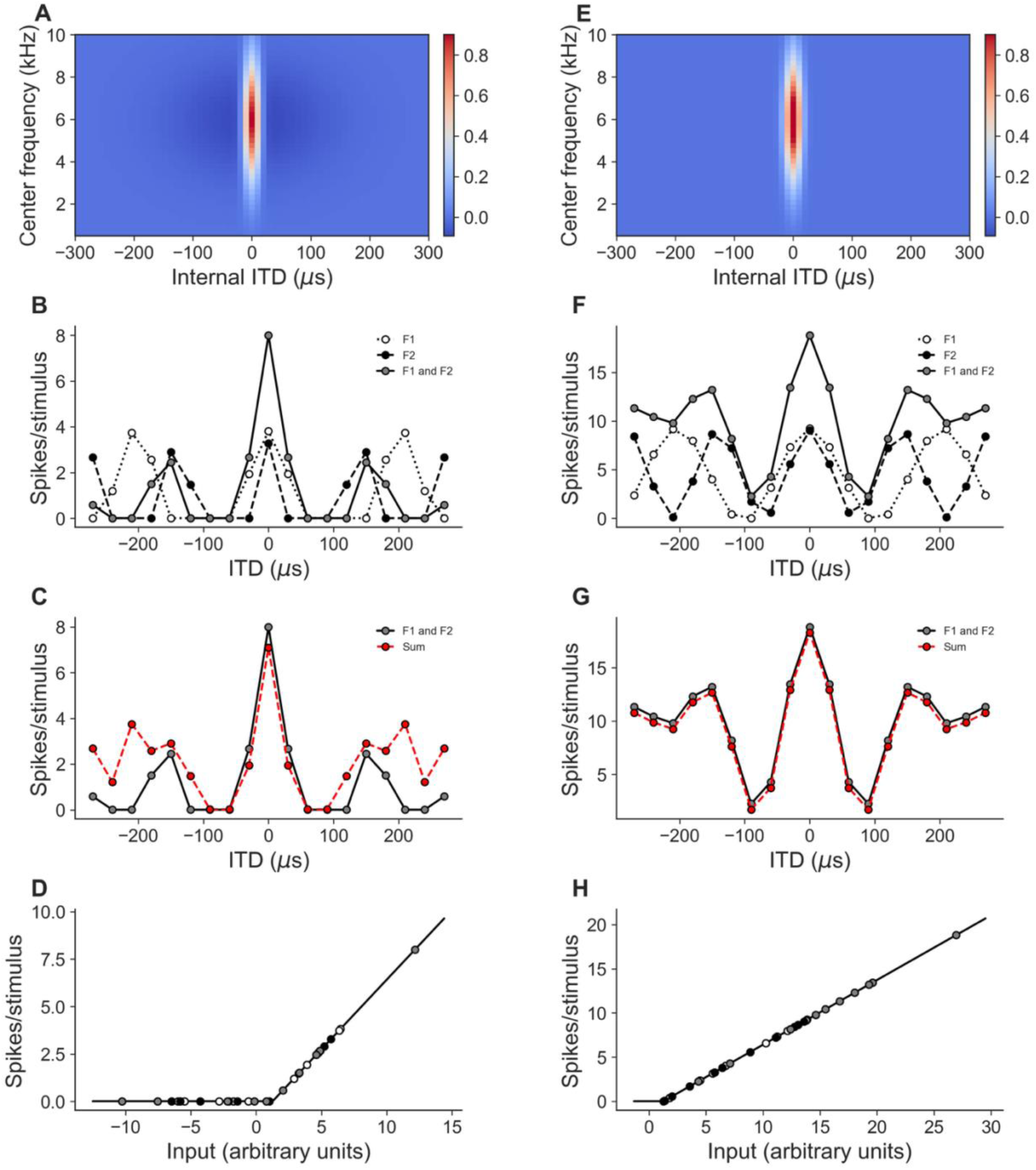
Nonlinear two-tone frequency integration produced by inhibition. (A-D) Responses with inhibitory surround weights. (A) Linear filter weight matrix with inhibitory surround. (B) ITD tuning curves for two tones at 6 and 8 kHz presented alone (F1, F2) and together (F1 and F2). (C) Comparison of the two-tone response (F1 andF2, black) with the sum of individual responses (Sum, red), showing facilitation at the main peak and suppression at side peaks. (D) Input-output function showing the nonlinear operating regime that produces facilitation and suppression. (E-H) Same neuron with inhibitory weights removed. (E) Filter weight matrix without surround inhibition. (F) ITD tuning curves for F1, F2, and F1 and F2 without inhibition. (G) The two-tone response now closely matches the arithmetic sum, indicating linear frequency integration. (H) Input-output function showing a linear operating regime. Removal of inhibition converts nonlinear to linear frequency integration, consistent with experimental observations of GABAergic blockade.

Nonlinear frequency integration that produces facilitation of the main peak can occur when the static nonlinearity is expansive. A rectified power-law static nonlinearity with a power greater than one can produce suppression at the side peaks where the input for one tone is inhibitory and facilitation at the main peak where the inputs from each tone produce a response on their own (Figure 7).

**Figure 7:**
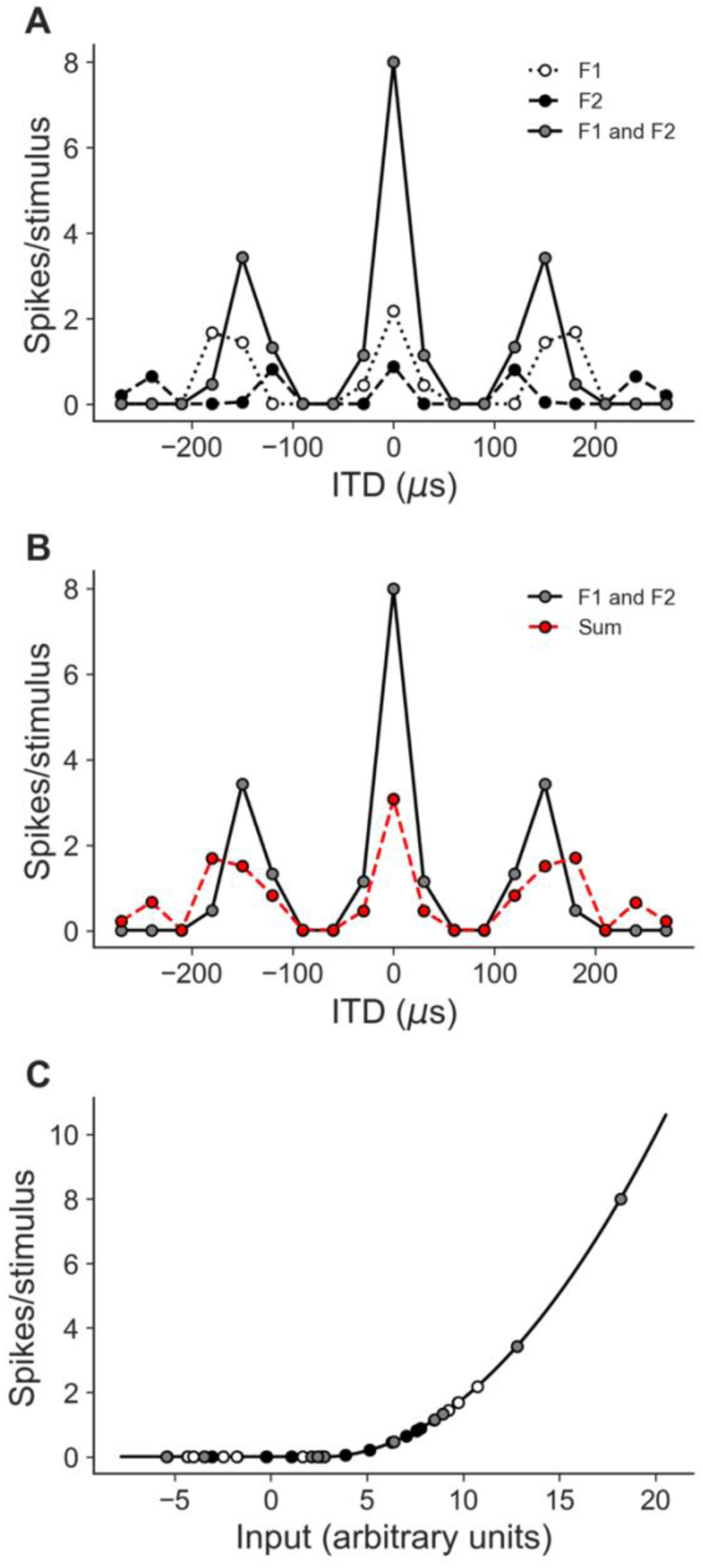
Nonlinear two-tone frequency integration with an expansive nonlinearity. (A) ITD tuning curves for two tones presented alone (F1, open circles; F2, filled circles) and together (F1 and F2, gray circles). The two-tone response shows strong main-peak facilitation and side-peak suppression. (B) Comparison of the two-tone response (F1 and F2, black) with the sum (Sum, red), demonstrating supralinear facilitation at the main peak and sublinear suppression at side peaks. (C) The expansive static nonlinearity (power > 1) that produces the nonlinear frequency integration. The operating point on the nonlinearity determines the degree of facilitation and suppression.

### Side peak suppression for broadband noise stimuli

ICx, and some ICcl neurons resolve phase ambiguity in ITD tuning by integrating inputs over frequency so that a larger response is produced at the main peak than at the side peaks of the ITD curve (Takahashi and Konishi, 1986; Mazer, 1998; Peña and Konishi, 2000). There is diversity across the population in how ITD tuning is shaped by frequency integration. In particular, the main peak and side peak of the ITD curve can respond differently to increases in the stimulus bandwidth (Mazer, 1998). Some of the diversity in responses can be explained by variation in the properties of the linear filter that describes linear subthreshold integration across frequency and ITD channels. The main peak of the ITD curve can show increasing, saturating, or non-monotonic responses to increasing stimulus bandwidth (Mazer, 1998). In the model, the main peak of the ITD curve occurs at the point where the “straight” part of the cross-correlation matrix is integrated over frequency. Increasing responses can be produced by having a linear filter with broad positive integration over the frequency axis (Figure 8AB). Saturating responses are produced when the linear filter is narrower along the frequency axis (Figure 8CD). Nonmonotonic responses that initially increase then decrease can be produced when the linear filter has a center-surround structure with positive weights near the best frequency and negative weights for frequencies (Figure 8EF).

**Figure 8:**
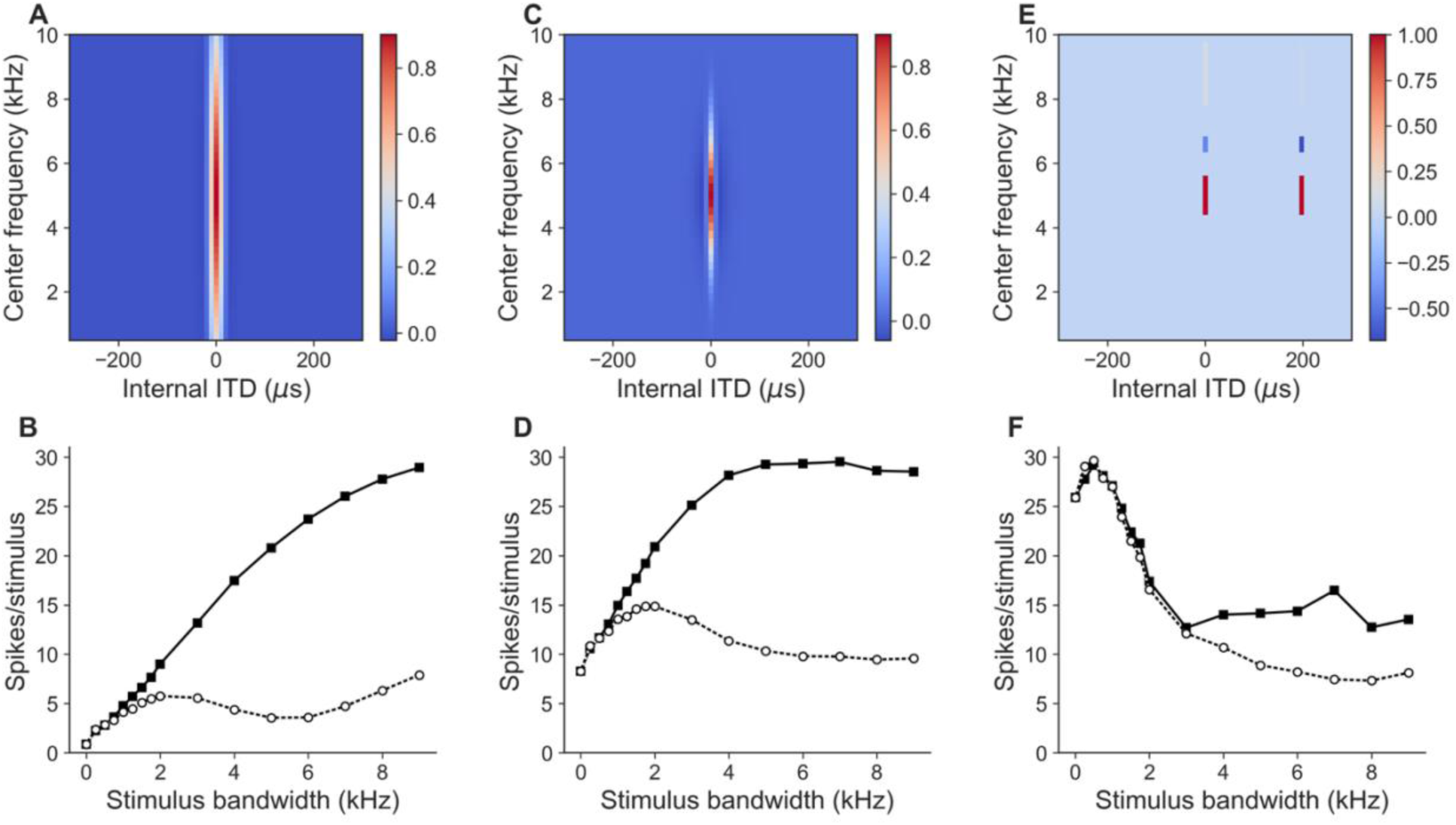
Side-peak suppression for broadband noise depends on linear filter shape. (A, B) Broad frequency filter. (A) Linear filter weight matrix with wide positive integration across the frequency axis. (B) Main-peak (filled squares) and side-peak (open circles) responses as a function of stimulus bandwidth, showing monotonically increasing responses at both peaks. (C, D) Narrow frequency filter. (C) Weight matrix with narrow integration across frequency. (D) The main peak saturates with increasing bandwidth while the side peak initially increases then decreases. (E, F) Bimodal frequency filter. (E) Weight matrix with the same weights at the best ITD and a phase equivalent ITD. (F) The main peak shows a non-monotonic response, increasing then decreasing with bandwidth, while main peak and side peak change nearly in parallel. These filter configurations reproduce the diversity of bandwidth-dependent responses observed experimentally.

The side peak of the ITD curve occurs where the cross-correlation peaks do not align across all frequencies. When the filter is narrow across frequency, the side peak will initially increase with stimulus bandwidth but then decrease as it incorporates more values at the trough of the cross-correlation (Figure 8CD). For wide frequency filters, the response at the side peak can again increase at high stimulus bandwidths where integration occurs over multiple peaks in the cross-correlation (Figure 8AB). An interesting type of response observed experimentally is where the main peak and side peak change nearly identically with stimulus bandwidth (Mazer, 1998). This can be produced in the model when the weights are similar at the internal ITD channels corresponding to the main peak and side peak (Figure 8EF).

### Population variation of side peak suppression and frequency tuning

A key challenge for any model of ITD processing in IC is explaining why SPS and frequency tuning width are uncorrelated across the neuronal population. In the standard cross-correlation model, neurons with broader frequency tuning naturally show greater SPS, but experimental data show that SPS and frequency half-width are uncorrelated (Mazer, 1998).

Figure 9 demonstrates that the modified model resolves this problem through independent variation of the linear filter bandwidth and static nonlinearity properties (Gorman et al., 2021). A population simulation varying the frequency filter width along with the threshold and power of the static nonlinearity reveals that SPS and frequency tuning half-width are uncorrelated across the population (Figure 9A-C). Different combinations of parameters produce different outcomes: neurons with broad frequency filters and a compressive nonlinearity show wide frequency tuning and low SPS; neurons with narrow frequency filters and an expansive nonlinearity show narrow frequency tuning and high SPS. The population scatter plot, overlaid with experimental data from Mazer (1998), shows that the model captures the observed uncorrelated relationship between SPS and frequency tuning width. Example model neurons demonstrate how the relationship between SPS and frequency tuning half-width can deviate from that predicted by the standard cross correlation model. The first example neuron has a broad frequency filter (Figure 9D) and a compressive nonlinearity (Figure 9E). The compressive nonlinearity reduces the SPS observed in the spiking output (Figure 9FG) and extends the broad frequency tuning created by the linear filter (Figure 9HI). The second example neuron has a narrow frequency filter (Figure 9J) and an expansive nonlinearity (Figure 9K). The expansive nonlinearity increases SPS and narrows the frequency tuning in the spiking output (Figure 9L-O). This demonstrates that the independent variation of nonlinearity shape, filter width, and threshold together can produce a more realistic model of ITD and frequency integration by IC neurons than the standard cross-correlation model.

**Figure 9:**
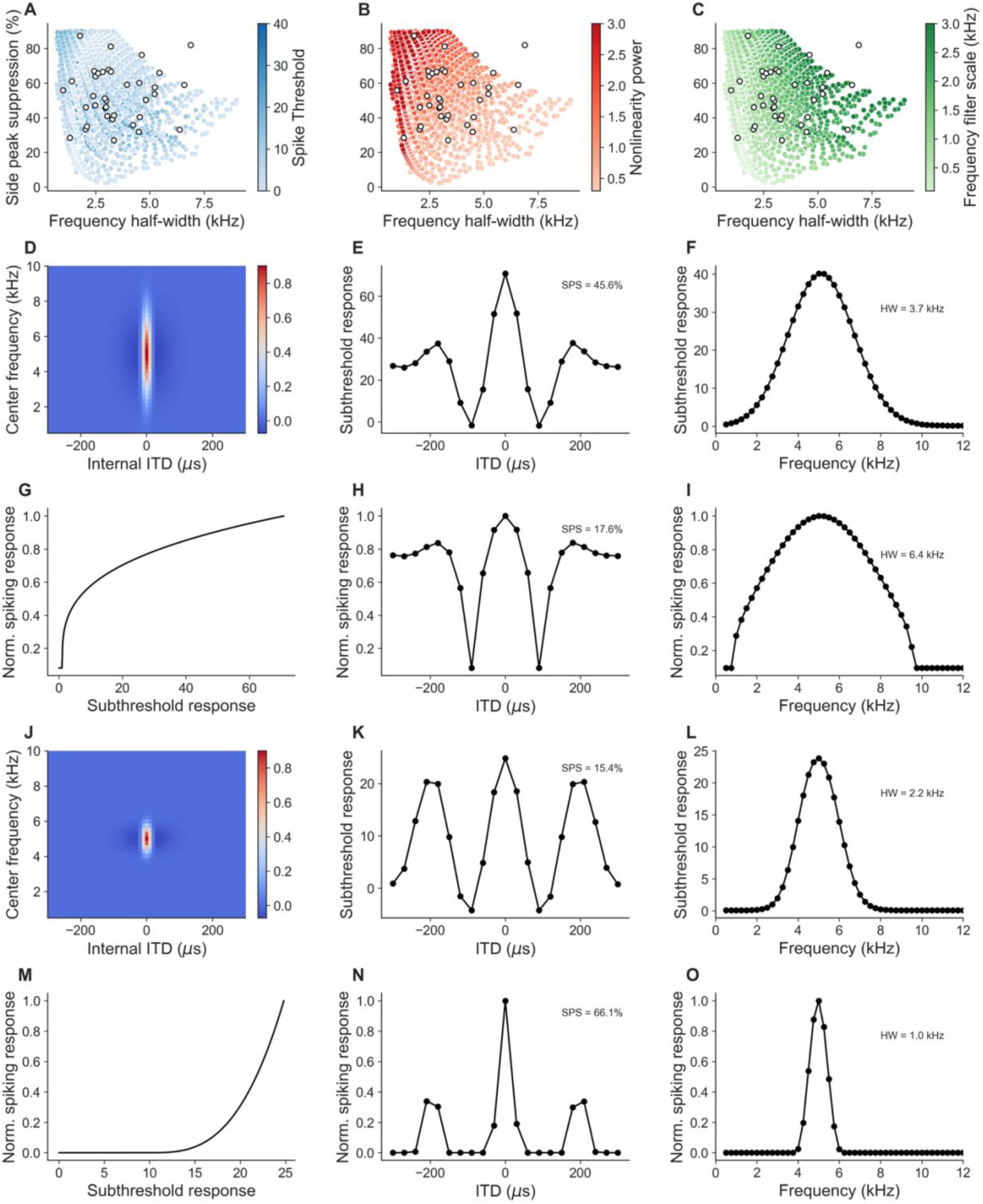
Independent variation of frequency tuning width and side-peak suppression across the model population. (A-C) Population scatter plots of SPS versus frequency half-width for model neurons with varying filter bandwidth, nonlinearity power, and threshold, overlaid with experimental data extracted from Figure 3 in Mazer (1998). The model reproduces the experimentally observed lack of correlation between SPS and frequency tuning width. (D-I) Example neuron with a broad frequency filter and compressive nonlinearity. (D) Wide linear filter weight matrix. (E) Compressive static nonlinearity. (F) ITD subthreshold response. (G) ITD tuning curve. (H) Frequency subthreshold response. (I) Frequency tuning curve. This combination produces wide frequency tuning and low SPS. (J-O) Example neuron with a narrow frequency filter and expansive nonlinearity. (J) Narrow linear filter weight matrix. (K) Expansive static nonlinearity. (L) ITD subthreshold response. (M) ITD tuning curve. (N) Frequency subthreshold response. (O) Frequency tuning curve. This combination produces narrow frequency tuning and high SPS, demonstrating that independent parameter variation decouples SPS from frequency tuning width.

### ITD tuning changes with ILD

Responses in ICx to combinations of ITD and ILD have been accurately described as a multiplication of one function of ITD and another function of ILD (Peña and Konishi, 2001, 2004). This shows that the shape of ITD tuning curves is largely independent of ILD in ICx. Multiplicative tuning to combinations of ITD and ILD emerges in ICcl, but responses are not uniformly multiplicative (Fischer et al., 2007). Specifically, ITD tuning can shift with ILD and the strength of ITD tuning can change with ILD, creating deviations from a purely multiplicative combination. Here, we show how the model can produce the observed responses.

### ITD shift with ILD

Shifts in best ITD with ILD occur in a subset of ICcl response (Fischer et al., 2007) (Figure 10). Shifting of best ITD with ILD is also observed in NL at the site of coincidence detection (Viete et al., 1997). The modified cross-correlation model produces shifts in best ITD with ILD by modifying the linear filter applied to the cross-correlation output dynamically based on an estimate of the ILD from the input signals. As in Fischer et al. (2008), the ILD is estimated from the gammatone filter outputs using a difference of the logarithms of the energy in the gammatone filter outputs. This estimate modifies the center frequency of the linear filter to produce a desired relationship between ILD and best ITD.

**Figure 10:**
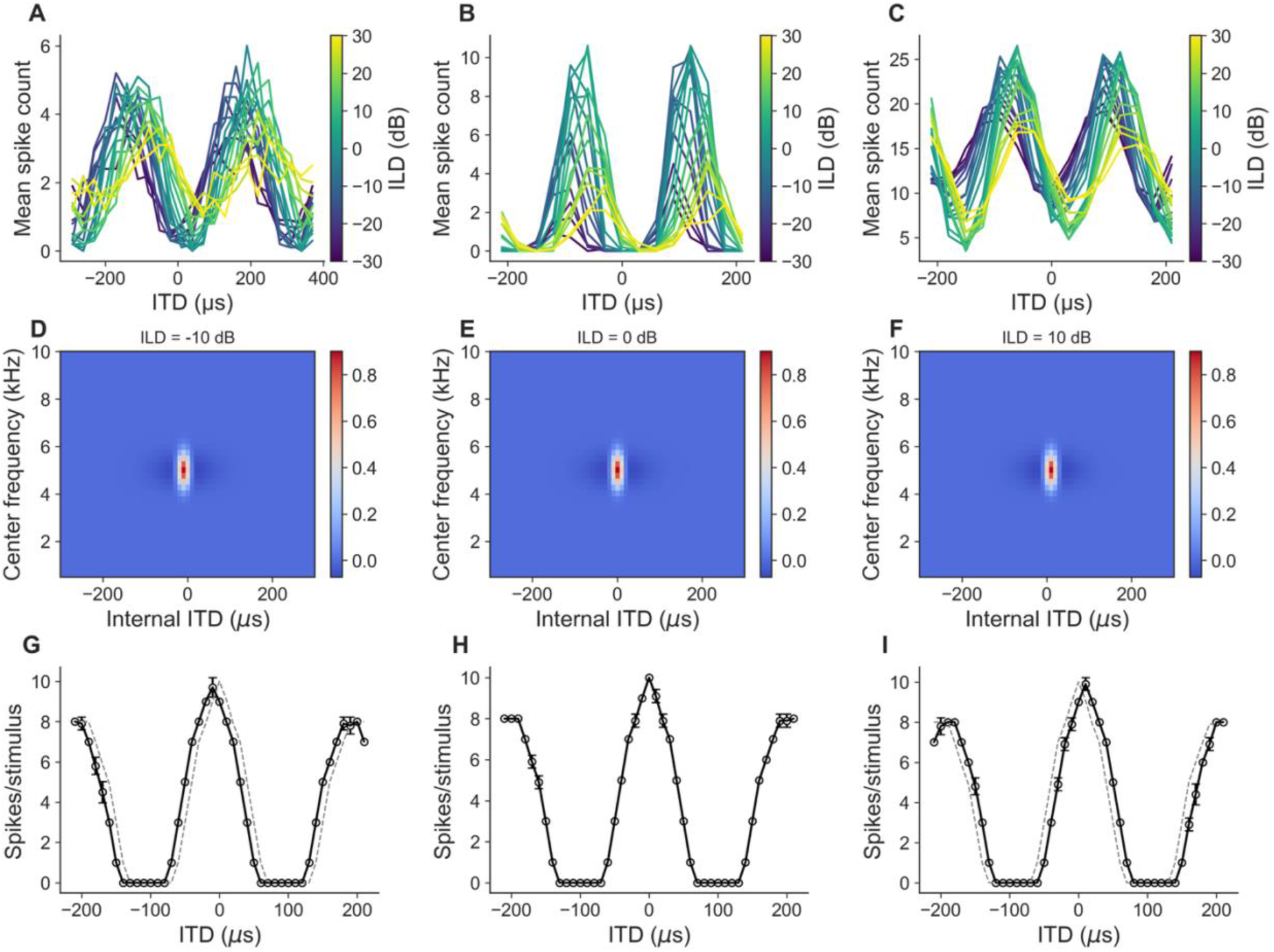
ILD-dependent shifts in best ITD. (A-C) Experimental ITD tuning curves from three ICcl neurons, with each curve colored by ILD (-30 to +30 dB). These neurons show systematic shifts in best ITD as a function of ILD. (D-F) Linear filter weight matrices of the model at three ILD values (-10, 0, and +10 dB), showing the ILD-dependent shift of the filter center along the internal ITD axis. (G-I) Corresponding model ITD tuning curves (solid lines with circles) at ILD = -10, 0, and +10 dB. Dashed gray curves show the reference tuning at ILD = 0 dB for comparison. The model produces the systematic shift in best ITD with ILD at a rate of 1.0 μs/dB.

Figure 10 (D-I) demonstrates systematic shifts in best ITD as a function of ILD. As ILD increases from -10 to 0 to +10 dB, the desired best ITD shifts systematically with a rate of 1.0 μs/dB. The filter weight matrices shift along the ITD axis with ILD, and the corresponding ITD tuning curves reflect this shift. This parameter can be tuned to match the magnitude of ITD shifts observed in individual neurons, allowing the model to capture the diversity of ILD-dependent best ITD shifts across the population.

### Loss of ITD tuning with ILD

In addition to shifts in best ITD with ILD, some ICcl neurons exhibit a decrease in the strength of ITD for some values of ILD (Fischer et al., 2007). As seen in Figure 11 A-C, the strength of ITD tuning can decrease while the neuron has a robust response to the sound.

**Figure 11:**
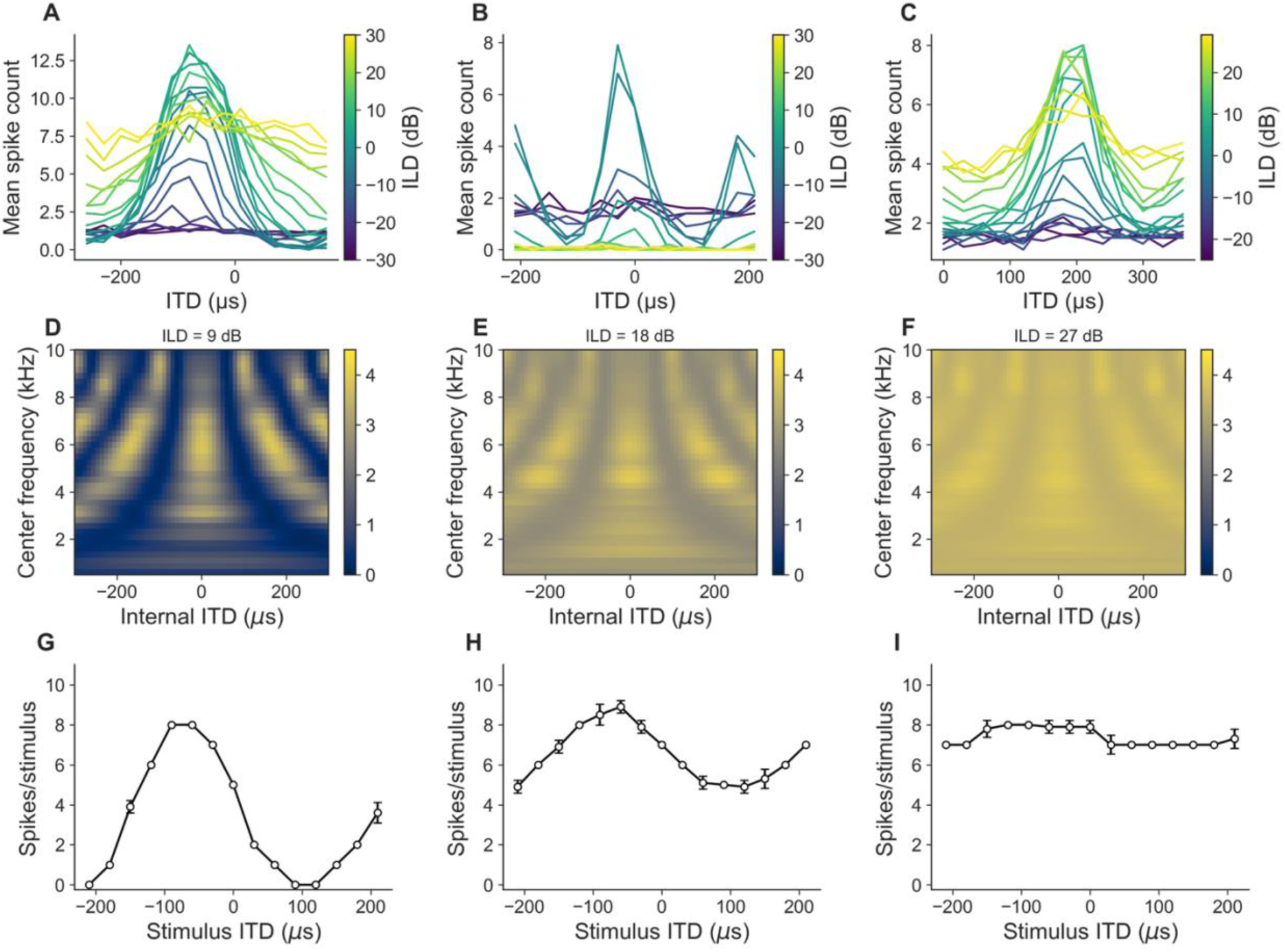
ILD-dependent loss of ITD tuning strength. (A-C) Experimental ITD tuning curves from three ICcl neurons, with each curve colored by ILD. These neurons show a progressive reduction in ITD tuning strength at extreme ILD values while maintaining an elevated firing rate. (D-F) Model linear filter weight matrices at ILD = 9, 18, and 27 dB. As ILD increases, the cross-correlation output is progressively replaced by a weighted average with a constant value, reducing ITD selectivity. (G-I) Corresponding model ITD tuning curves at ILD = 9 dB (strong ITD tuning), 18 dB (moderate ITD tuning), and 27 dB (weak ITD tuning). The model maintains an elevated response across ITD values while the peak-trough modulation depth decreases, reproducing the experimental observation that some ICcl neurons lose ITD selectivity at certain ILD values without losing responsiveness.

The modified cross-correlation model produces changes in the strength of ITD tuning with ILD by replacing the output of the cross-correlation with a weighted average of the cross-correlation and a constant value, where the relative weights are determined by the estimated ILD. This allows the peak-trough difference of the ITD tuning curve to depend on ILD. Figure 11 D-I demonstrates this mechanism where ITD tuning is strong at ILD = 9 dB, moderate at ILD = 18, and weak at ILD = 27 dB, while firing rate remains elevated from baseline across ILD values. This matches the experimental observation that some neurons maintain responsiveness across ILD values while losing their ITD selectivity.

The model demonstrates that these two phenomena, ITD shift and loss of vector strength, can be controlled independently through separate parameters, allowing flexible capture of the diversity of ILD-dependent ITD modulation observed in the population.

### Parameter selection

A complete characterization of the model requires determining the parameters that control ITD tuning, frequency integration, rate-level functions, and ILD dependence. Several parameters can be computed directly from desired response properties. Other parameters are determined using SBI, a machine learning approach that efficiently identifies parameter combinations producing desired outputs.

Figure 12 demonstrates population generation using SBI-based parameter fitting. For each neuron, SBI was used to find gain control power 𝛼 and bias 𝛾 values producing a desired rate-level function shape, and to find linear filter width and static nonlinearity power producing desired SPS and frequency half-width values. There is good agreement between desired and measured properties across the population: best frequency, frequency half-width, best ITD, SPS, ITD shift rate with ILD, and ITD dynamic range at extreme ILD values are all highly correlated with the desired values (Figure 12 A-F). The desired rate-level functions of the linear input to the neuron are specified using a form that describes rate-level functions of auditory-nerve fibers (methods) (Köppl and Yates, 1999). The desired minimum and maximum firing rates are highly correlated with the measured values (Figure 12G, H). The shape parameters 𝐴_2_, 𝐴_3_, and 𝐴_4_ are also correlated with the measured values, but less so than the minimum and maximum firing rates (Figure 12 I-K). However, combinations of these parameters produce similar rate-level functions, so they are not as accurately determined by the model fitting procedure. Nevertheless, the relative error between the desired and measured rate-level function is small (Figure 12L).

**Figure 12:**
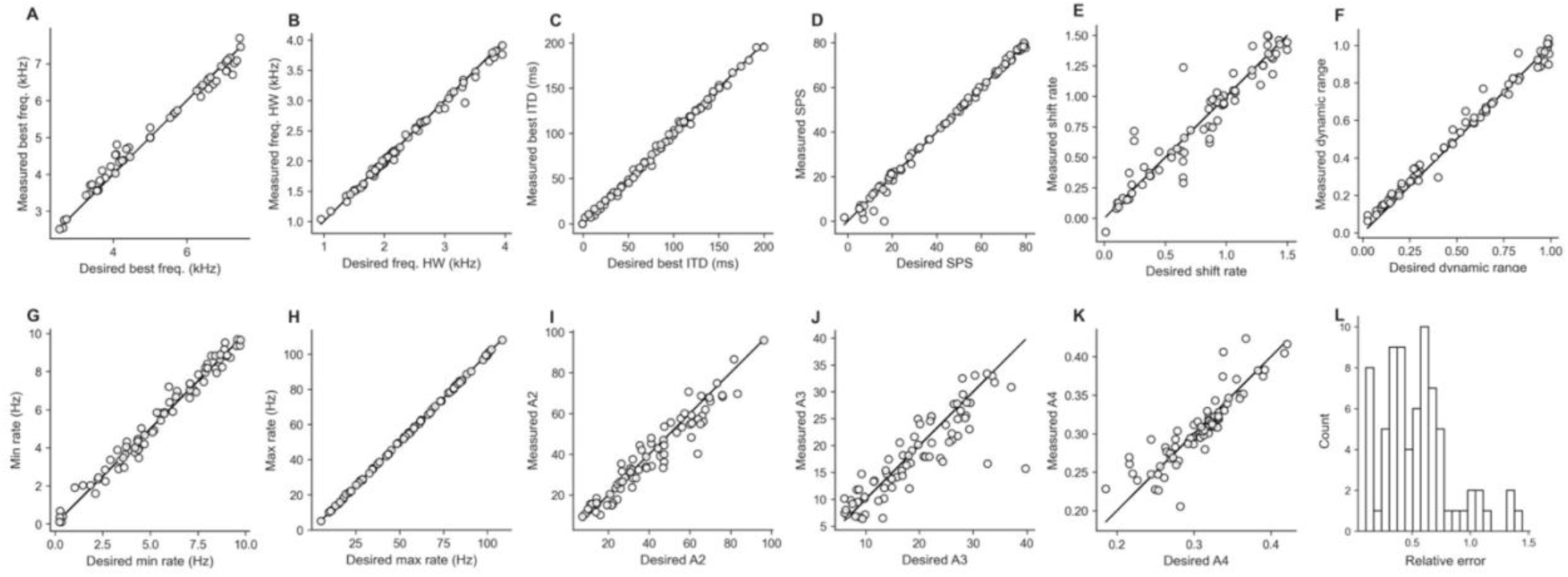
Population parameter fitting using simulation-based inference. Validation of parameter determination for a population of model IC neurons. Scatter plots show desired versus measured values for each response property: (A) best frequency, (B) frequency half-width, (C) best ITD, (D) side-peak suppression, (E) ITD shift rate with ILD, (F) ITD dynamic range at ILD = 30 dB, parameters of the rate-level function (G-K). (L) Relative error between the desired and measured rate-level functions. The close agreement across all properties demonstrates that SBI can efficiently identify parameter combinations that produce target response characteristics, enabling generation of realistic model populations with diverse ITD tuning across frequency channels.

This demonstrates that the model fitting approach using SBI can efficiently determine parameters to match desired response properties, and that the model can generate realistic populations of IC neurons with diverse ITD selectivity across frequency channels. The parameter flexibility and analytical tractability of the model make it suitable for exploring population coding strategies in the owl’s auditory midbrain.

## Discussion

The cross-correlation model has long served as a foundational framework for describing ITD tuning in the barn owl’s auditory system. It has been remarkably successful in explaining a variety of auditory responses, particularly those observed at the site of coincidence detection in NL and in IC. The standard model, characterized by linear filtering and summation of cross-correlated signals, can reproduce sinusoidal ITD tuning curves and frequency integration properties seen in a subset of neurons. However, the linear version of the model is fundamentally limited in its ability to capture the full diversity of ITD tuning observed in IC neurons. Many neurons show response characteristics that deviate from strict linear predictions, including nonlinear frequency interactions, threshold effects, changes in best ITD with ILD, and varied SPS versus frequency tuning width relationships.

The work presented here is not unique in modifying the cross-correlation model to describe aspects of ITD processing. Previous work has introduced modifications to the cross-correlation model to describe IC responses (Albeck and Konishi, 1995; Keller and Takahashi, 2005; Fischer et al., 2009). In this study, we extend the cross-correlation framework to include flexible, parameterizable linear filters and nonlinearities. This expanded model architecture enables us to demonstrate how specific filter shapes and nonlinear transformations can generate a wide range of realistic responses to tones and broadband noise across the IC.

We have presented a modified cross-correlation model of ITD computation in the barn owl’s IC that successfully addresses the major discrepancies between the standard cross-correlation model and experimental observations. The key modifications, gain control for stimulus level dependence, ILD-dependent filter structure, frequency-dependent weighting via the linear filter, and nonlinear transformation, work together to produce the response diversity observed in ICcl and ICx neurons while maintaining a framework grounded in the classical Jeffress model of ITD coding. This more flexible and physiologically grounded version of the cross-correlation model provides a foundation for accurate modeling of population coding in the barn owl’s midbrain. By allowing variation in linear filter structure, gain control mechanisms, and spiking nonlinearities, the model captures both the common principles and the diversity of auditory spatial tuning across IC subnuclei. These features are critical for understanding how the auditory system encodes sound source location under naturalistic conditions, where stimulus features vary widely in time, frequency, and amplitude.

The use of simulation-based inference for parameter fitting represents a practical approach to identifying biophysically plausible model parameters when experimental constraints are numerous and not easily relatable to parameters. The close agreement between desired and measured population response properties (Figure 12) validates the model’s capacity to serve as a tool for systematic exploration of population coding. Future studies can use this approach to investigate how constraints on population architecture, such as the range of frequency tuning widths or the distribution of nonlinearity powers, shape auditory space coding and behavioral sound localization.

The barn owl’s ITD circuitry remains a powerful paradigm for studying neural computation. The cross-correlation model continues to be a central tool in interpreting these circuits, and our work enhances its descriptive power. However, several limitations and future directions remain. The model presently lacks explicit representation of temporal dynamics of inhibition, simplifying it to static operations. The model does not include a full description of ILD tuning in IC, which is also varied across ICcl and ICx (Moiseff, 1989; Adolphs, 1993b; Fischer et al., 2007). Additionally, the variability of spiking responses has not been modeled (Beckert et al., 2020). While there remain elements of ICcl and ICx responses to include in the model, the model presented here captures the diversity of ITD-dependent responses in IC.

In summary, a cross-correlation framework, augmented with biologically inspired flexible nonlinearities and linear filters, can capture the rich diversity of ITD tuning observed in the owl’s auditory midbrain. These features are critical for understanding how the auditory system encodes sound source location under naturalistic conditions, where stimulus features vary widely in time, frequency, and amplitude. This provides a powerful platform for investigating population coding and sensory representation in neural circuits that have been evolutionarily optimized for precise spatial hearing.

## Acknowledgements

This work was funded by the National Institute of Neurological Disorders and Stroke (1RF1NS132812-01)

## Notes

https://github.com/brian-fischer/Flexible-xcorr-model

### Competing Interest Statement

The authors have declared no competing interest.

## References

Adolphs R (1993a) Acetylcholinesterase staining differentiates functionally distinct auditory pathways in the barn owl. J Comp Neurol 329:365–377.

Adolphs R (1993b) Bilateral inhibition generates neuronal responses tuned to interaural level differences in the auditory brainstem of the barn owl. J Neurosci 13:3647–3668.

Albeck Y, Konishi M (1995) Responses of neurons in the auditory pathway of the barn owl to partially correlated binaural signals. J Neurophysiol 74:1689–1700.

Arthur BJ (2004) Sensitivity to spectral interaural intensity difference cues in space-specific neurons of the barn owl. J Comp Physiol A 190:91–104.

Beckert MV, Fischer BJ, Pena JL (2020) Effect of stimulus-dependent spike timing on population coding of sound location in the owl’s auditory midbrain. eNeuro 7.

Brainard MS, Knudsen EI, Esterly SD (1992) Neural derivation of sound source location: resolution of spatial ambiguities in binaural cues. J Acoust Soc Am 91:1015–1027.

Brette R, Gerstner W (2005) Adaptive exponentialiIntegrate-and-fire model as an effective description of neuronal activity. J Neurophysiol 94:3637–3642.

Carr CE, Konishi M (1988) Axonal delay lines for time measurement in the owl’s brainstem. Proc Natl Acad Sci 85:8311–8315.

Cazettes F, Fischer BJ, Pena JL (2014) Spatial cue reliability drives frequency tuning in the barn Owl’s midbrain. eLife 3:e04854.

Cranmer K, Brehmer J, Louppe G (2020) The frontier of simulation-based inference. Proc Natl Acad Sci 117:30055–30062.

Euston DR, Takahashi TT (2002) From spectrum to space: the contribution of level difference cues to spatial receptive fields in the barn owl inferior colliculus. J Neurosci 22:284–293.

Ferger R, Shadron K, Fischer BJ, Peña JL (2021) Barn owl’s auditory space map activity matching conditions for a population vector readout to drive adaptive sound-localizing behavior. J Neurosci 41:10305–10315.

Fischer BJ, Anderson CH, Peña JL (2009) Multiplicative auditory spatial receptive fields created by a hierarchy of population codes. PloS One 4:e8015.

Fischer BJ, Christianson GB, Peña JL (2008) Cross-correlation in the auditory coincidence detectors of owls. J Neurosci 28:8107–8115.

Fischer BJ, Peña JL (2009) Bilateral matching of frequency tuning in neural cross-correlators of the owl. Biol Cybern 100:521–531.

Fischer BJ, Peña JL (2011) Owl’s behavior and neural representation predicted by Bayesian inference. Nat Neurosci 14:1061–1066.

Fischer BJ, Peña JL, Konishi M (2007) Emergence of multiplicative auditory responses in the midbrain of the barn owl. J Neurophysiol 98:1181–1193.

Fontaine B, Peña JL, Brette R (2014) Spike-threshold adaptation predicted by membrane potential dynamics in vivo. PLoS Comput Biol 10:e1003560.

Fujita I, Konishi M (1991) The role of GABAergic inhibition in processing of interaural time difference in the owl’s auditory system. J Neurosci 11:722–739.

Gold JI, Knudsen EI (2000) Abnormal auditory experience induces frequency-specific adjustments in unit tuning for binaural localization cues in the optic tectum of juvenile owls. J Neurosci 20:862–877.

Gorman JC, Tufte OL, Miller AVR, DeBello WM, Peña JL, Fischer BJ (2021) Diverse processing underlying frequency integration in midbrain neurons of barn owls. PLOS Comput Biol 17:e1009569.

Jeffress LA (1948) A place theory of sound localization. J Comp Physiol Psychol 41:35–39.

Keller CH, Takahashi TT (1996) Binaural cross-correlation predicts the responses of neurons in the owl’s auditory space map under conditions simulating summing localization. J Neurosci 16:4300–4309.

Keller CH, Takahashi TT (2005) Localization and identification of concurrent sounds in the owl’s auditory space map. J Neurosci 25:10446–10461.

Knudsen EI, Konishi M (1978) A neural map of auditory space in the owl. Science 200:795–797.

Knudsen EI, Konishi M, Pettigrew JD (1977) Receptive fields of auditory neurons in the owl. Science 198:1278–1280.

Konishi M (1991) Deciphering the brain’s codes. Neural Comput 3:1–18.

Köppl C (1997) Frequency tuning and spontaneous activity in the auditory nerve and cochlear nucleus magnocellularis of the barn owl Tyto alba. J Neurophysiol 77:364–377.

Köppl C, Yates G (1999) Coding of sound pressure level in the barn owl’s auditory nerve. J Neurosci 19:9674–9686.

Mazer JA (1998) How the owl resolves auditory coding ambiguity. Proc Natl Acad Sci 95:10932–10937.

Moiseff A (1989) Bi-coordinate sound localization by the barn owl. J Comp Physiol A Neuroethol Sens Neural Behav Physiol 164:637–644.

Mori K (1997) Across-frequency nonlinear inhibition by GABA in processing of interaural time difference. Hear Res 111:22–30.

Peña JL, Konishi M (2000) Cellular mechanisms for resolving phase ambiguity in the owl’s inferior colliculus. Proc Natl Acad Sci 97:11787–11792.

Peña JL, Konishi M (2001) Auditory spatial receptive fields created by multiplication. Science 292:249–252.

Peña JL, Konishi M (2002) From postsynaptic potentials to spikes in the genesis of auditory spatial receptive fields. J Neurosci 22:5652–5658.

Peña JL, Konishi M (2004) Robustness of multiplicative processes in auditory spatial tuning. J Neurosci Off J Soc Neurosci 24:8907–8910.

Peña JL, Viete S, Albeck Y, Konishi M (1996) Tolerance to sound intensity of binaural coincidence detection in the nucleus laminaris of the owl. J Neurosci 16:7046–7054.

Takahashi T, Konishi M (1986) Selectivity for interaural time difference in the owl’s midbrain. J Neurosci 6:3413–3422.

Takahashi TT, Konishi M (1988) Projections of the cochlear nuclei and nucleus laminaris to the inferior colliculus of the barn owl. J Comp Neurol 274:190–211.

Tejero-Cantero A, Boelts J, Deistler M, Lueckmann J-M, Durkan C, Gonçalves PJ, Greenberg DS, Macke JH (2020) sbi: A toolkit for simulation-based inference. J Open Source Softw 5:2505.

Viete S, Peña JL, Konishi M (1997) Effects of interaural intensity difference on the processing of interaural time difference in the owl’s nucleus laminaris. J Neurosci 17:1815–1824.

Wagner H, Mazer JA, von Campenhausen M (2002) Response properties of neurons in the core of the central nucleus of the inferior colliculus of the barn owl. Eur J Neurosci 15:1343–1352.

Wagner H, Takahashi T, Konishi M (1987) Representation of interaural time difference in the central nucleus of the barn owl’s inferior colliculus. J Neurosci 7:3105–3116.

